# Sex-dependent transcriptional control of cardiac electrophysiology by histone acetylation modifiers based on the GTEx database

**DOI:** 10.1101/2022.04.19.488752

**Authors:** Michael P. Pressler, Anelia Horvath, Emilia Entcheva

## Abstract

Development of safer drugs based on epigenetic modifiers, e.g. histone deacetylase inhibitors (HDACi), requires better understanding of their effects on cardiac electrophysiology. Using RNAseq data from the genotype-tissue-expression database (GTEx), we created models that link the abundance of chromatin modifiers, such as histone acetylation enzymes (HDAC/SIRT/HATs), and the gene expression of ion channels (IC) via select cardiac transcription factors (TFs) in male and female adult human hearts (left ventricle, LV). Gene expression data (transcripts per million, TPM) from GTEx donors (21 to 70 y.o.) were filtered, normalized and transformed to Euclidian space to allow quantitative comparisons in 84 female and 158 male LVs. Sex-specific partial least-square (PLS) regression models, linking gene expression data for HDAC/SIRT/HATs to TFs and to ICs gene expression, revealed tight co-regulation of cardiac ion channels by HDAC/SIRT/HATs, with stronger clustering in the male LV. Co-regulation of genes encoding excitatory and inhibitory processes in cardiac tissue by the histone modifiers may help their predominantly net-neutral effects on cardiac electrophysiology. *ATP1A1*, encoding for the Na/K pump, represented an outlier - with orthogonal regulation by the histone modifiers to most of the ICs. The HDAC/SIRT/HAT effects were mediated by strong (+) TF regulators of ICs, e.g. *MEF2A* and *TBX5*, in both sexes. Furthermore, for male hearts, PLS models revealed a stronger (+)/(-) mediatory role on ICs for *NKX25* and *TGF1B/KLF4*, respectively, while *RUNX1* exhibited larger (-) TF effects on ICs in females. Male-trained PLS models of HDAC/SIRT/HAT effects on ICs underestimated the effects on some ICs in females. Insights from the GTEx dataset about the co-expression and transcriptional co-regulation of histone-modifying enzymes, transcription factors and key cardiac ion channels in a sex-specific manner can help inform safer drug design.

## Introduction

Over the last decade, large-scale annotated biomedical data collections, have been assembled, such as the genotype-tissue expression (GTEx) dataset[1-3], combining RNAseq and other technologies to enable discovery of molecular mechanisms of human disease with unprecedented throughput. Transcriptomics data can be linked to subject’s demographics and other characteristics, and molecular relationships can be parsed by sex, age, tissue context and other characteristics. Now, these resources are being leveraged to estimate safety risk for drugs and to predict the outcomes of clinical trials[4-6], in hopes to reduce cost, time for drug development and potential side effects. Such analyses using the GTEx data have brought awareness to tissue specificity[1], cell specificity[7] and the impact of sex on key molecular pathways and transcriptional control mechanisms[7-10]. For example, the GTEx dataset has been useful in identifying heart-specific genes implicated in pathologies, and in exploring cardiac tissue-specific relationships and regulators as well as transcriptome-wide differences between the ventricles and the atria [8] [9, 11-16]. Such unique cardiac targets can be useful in designing new and safer therapies. Furthermore, transcriptional principles of ion channel co-expression, underlying bioelectrical stability (lower risk of early afterdepolarizations and other arrhythmias), have been uncovered using the GTEx data from the left ventricle (LV) and validated in vitro using human induced pluripotent stem-cell-derived cardiomyocytes, iPSC-CMs[14].

Avoiding cardiotoxicity is a critical consideration in drug design. Cardiotoxicity is at least in part due to side effects of drugs on key cardiac ion channels (directly modulating their function, affecting their transcription or trafficking), thus leading to increased arrhythmia risk[17]. Prior failed drugs and recent computational data stress the need for addressing diversity in responses – for example, a drug-response risk classifier trained on male data underpredicts arrhythmia risk in females[18]. Hormonal effects are now believed to only partially explain sex differences in these functional responses, as the GTEx data analysis has revealed that about 37% of all genes (majority of these autosomal) exhibit sex-differential gene expression in at least one tissue, including in the LV of the heart, where over 1300 genes have differential sex expression[7]. Importantly, among the most sex-differentially expressed genes are those involved in ion transport processes and response to drugs, including ion channels and CYP enzymes[6, 7, 19]. Furthermore, sex differences in gene expression in the heart[8, 9, 20] include differences in the expression of epigenetic modifiers, in chromatin accessibility and the related regulatory networks[21, 22]. Even when sex differences in transcription are not dramatic, the cumulative effect of these at the system level can be functionally impactful because of the differential effects on multiple key regulators, including cardiac transcription factors[10].

Here we consider the relationship between a class of epigenetic regulators – histone acetylation modifiers – and key cardiac ion channels, defining excitation, repolarization, calcium handling and cell-cell coupling in the heart’s LV. These include histone deacetylases (HDACs and sirtuins, SIRTS) acting as “acetylation erasers” and histone acetyltransferases (HATs) acting as “writers”, which operate to balance chromatin acetylation state and accessibility for transcription factors to regulate gene expression[23] (**Figure 1**). Sirtuins, SIRTs, represent a sub-class of HDACs with NAD+ dependence, therefore, linking metabolism to post-translational modifications. In general, decreased action of HDACs and SIRTs tilts the balance to increased acetylation, a more open chromatin state and increase in transcription of certain genes (apoptosis and other growth control processes being among the most affected ones), of particular importance in anti-cancer strategies. HDAC inhibitors (HDACi) have emerged in the last two decades as a novel anti-cancer therapy, with four of them FDA approved and several hundred clinical trials on the way[24, 25]. Early analysis of safety (from clinical trials)[26, 27] and in vitro work with human iPSC-CMs[28-30] indicates that overall HDACi are relatively safe, with few adverse events, including arrhythmias, more prevalent for non-selective HDACi, e.g. panobinostat and vorinostat, and less prevalent for selective inhibitors, such as entinostat. Therefore, the identity of the HDACis and their specific targets among the cardiac electrophysiology key players are of high importance in achieving safer therapeutics. Beyond initial animal studies that have established select engagement of cardiac transcription factors (TFs) upon HDACi-related chromatin loosening and their role in controlling ion channel (IC) gene expression in the heart[30, 31], a more comprehensive model linking HDACs/SIRTs/HATs to control of ion channel transcription in the human heart via the action of cardiac TFs[31-33] is highly desirable.

**Figure 1:**
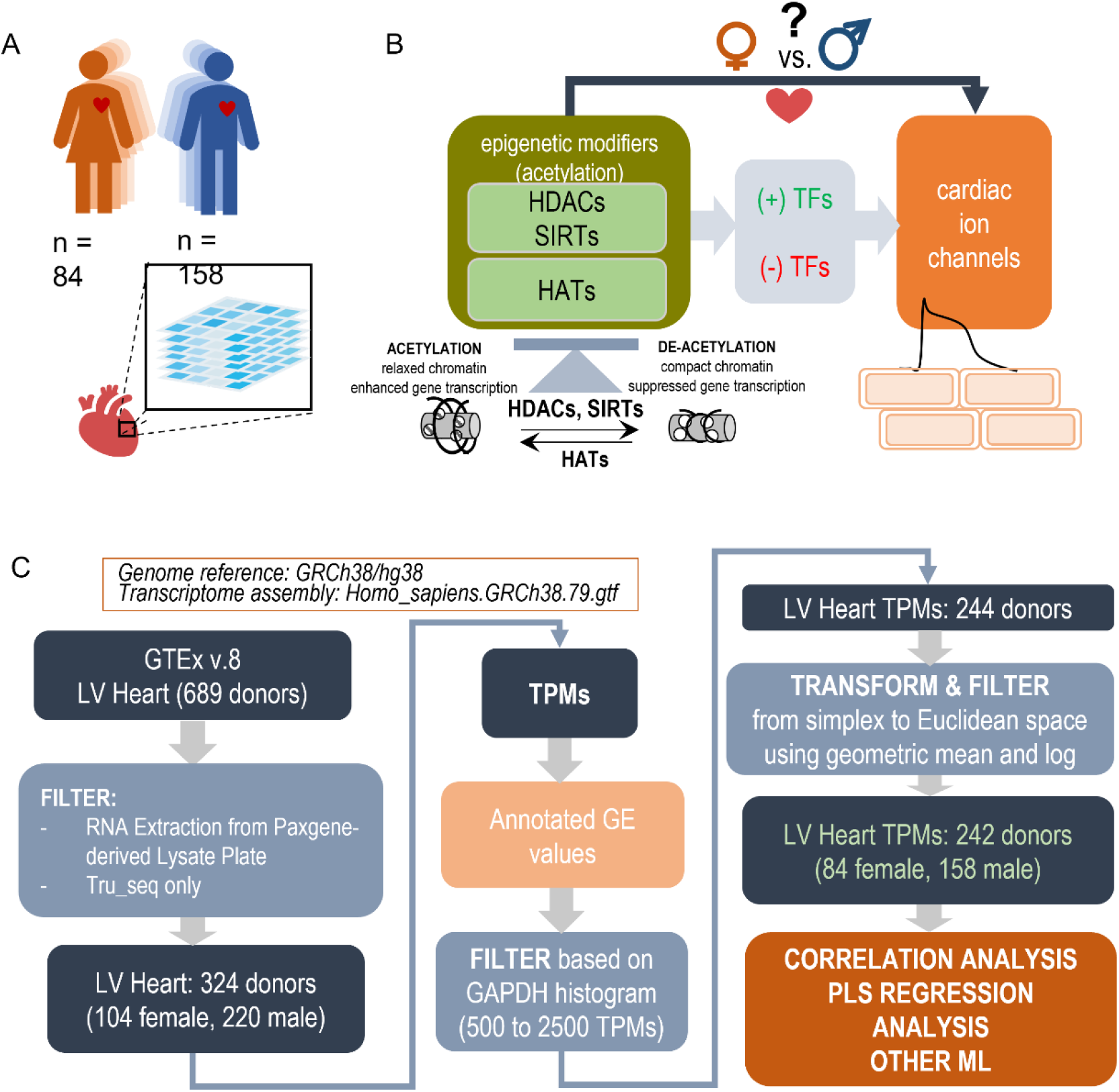
Study design outline and processing steps: Using the GTEx database to inform PLS regression models of the effects of histone modifiers (HDACs, SIRTs, HATs) on cardiac ion channels via key TFs in the adult male and female left ventricle (LV). A. LV bulk RNAseq data from 84 female and 158 male adults were used from the GTEx dataset. B. The performed analysis examines how the balance of HDACs, SIRTs, and HATs may affect cardiac ion channel transcription in a sex-dependent manner. HDACs and SIRTs counter the action of HATs, which acetylate chromatin thus increasing its accessibility for key cardiac TFs to act on genes of interest. HDACs, SIRTs, and HATs negatively or positively regulate the effects of TFs and consequently change the expression of cardiac ion channels. C. Processing pipeline: Starting with 689 donors in the GTEx v.8 dataset, after filtering and transformations, 242 donors were used for correlation and PLS regression analysis. The processing steps include TPM filtering based on GAPDH levels to yield similar normal distributions and a log-ratio transformation from simplex to Euclidian space.

Our objective was to use the GTEx RNAseq dataset to construct simplified computational models linking these classes of gene regulators, and to investigate if sex-specific differences exist. We chose partial least-square (PLS) regression modeling[34-36] to quantify the relationships in female and male hearts. Projection of the many master regulators into a reduced-dimension latent structures through PLS helps to elucidate co-regulation mechanisms of cardiac electrophysiology by these histone modifiers and to identify potential key mediators among the cardiac TFs. The exponential rise of publicly accessible human transcriptomics datasets can yield continuous refinement of such models for better stratification of drug action. Such models can help understand the specific effects of histone acetylation enzymes on the heart and guide the design of safer anti-cancer therapeutics, minimizing cardiac effects, or help targeted HDACi design for cardiac use to purposefully augment cardiac function.

## Methods

### Classes of human genes in the models

The human genes analyzed here are listed in **Table 1**. These included the four classes of histone deacetylases, HDACs, for a total of 18: class I (*HDAC1, 2, 3 and 8*), class IIa (*HDAC4, 5, 7, 9*), class IIb (*HDAC6 and 10*), class III (*SIRT1 to 7*) and class IV (*HDAC11*); 15 key histone acetyltransferases HATs, including EP300; 19 cardiac transcription factors (TFs) and 10 genes encoding proteins that form key cardiac ion channels, ICs.

**Table 1.** Genes considered in the models.

|  |  |
| --- | --- |
| **HDACs (11)** | *HDAC1, HDAC2, HDAC3, HDAC4, HDAC5, HDAC6, HDAC7, HDAC8, HDAC9, HDAC10, HDAC11* |
| **SIRTs (7)** | *SIRT1, SIRT2, SIRT3, SIRT4, SIRT5, SIRT6, SIRT7* |
| **HATs (15)** | *KAT2A, KAT2B, HAT1, ATF2, KAT5, KAT6A, KAT6B, KAT7, EP300, CREBBP, NCOA1, NCOA3, TAF1, GTF3C1, CLOCK* |
| **TFs (19)** | *FOXO1, FOXO3, GATA4, GATA6, HIF1A, KLF4, KLF5, MEF2A, NFAT5, NFKB1, NKX25, NOTCH1, RUNX1, SHMT2, SOD1, TBX5, TGFB1, TRIM28, YY1* |
| **ICs (10)** | *SCN5A, CACNA1C, KCNH2, KCNQ1, KCNJ2, ATP1A1, SLC8A1, ATP2A2, RYR2, GJA1* |

### Sourcing the data from GTEx v.8

The analyses are based on the use of study data obtained from the dbGaP web site, under dbGaP accession number phs000424.v8.p2 (GTEx, https://gtexportal.org/home/datasets). Expression levels in transcripts per million (TPM) derived from RNA-seq data were downloaded on 08/28/2020 as a single file combining all body sites and cell types from the GTEx database. From this file, 324 TPM datasets of samples (103 female and 220 male samples) obtained from heart left ventricle (LV) were extracted based on accession numbers listed in the de-identified, open access version of the sample annotations available in dbGaP (GTEx_Analysis_v8_Annotations_SampleAttributesDS, downloaded on 08/28/2020, https://gtexportal.org/home/datasets). The subjects age, gender and additional phenotype features were obtained from the de-identified, open access version of the subject phenotypes available in dbGaP (GTEx_Analysis_v8_Annotations_SubjectPhenotypesDS, downloaded on 08/28/2020, https://gtexportal.org/home/datasets). Subjects range in age between 20 and 71 y.o. These RNA-seq libraries were generated using non-strand specific, polyA-based Illumina TruSeq protocol and sequenced to a median depth of 78 million 76-bp paired-end reads. Annotated gene expression was obtained using genome reference Homo_sapiens.GRCh38.79.gtf.

### Data preprocessing

As indicated in **Figure 1**, TPM values were filtered, normalized and transformed before analysis. The large scale of the GTEx project and the span over a decade implies continued optimization of the sequencing technology and sample handling. Thus, the dataset contains samples of variable quality, mostly influenced by the timing of death with respect to sample processing[37]. In addition to the quality control applied by the consortium, we chose to filter the data by *GAPDH* levels (leaving samples with 500 to 2500 TPMs in *GAPDH*)- a prominently expressed gene that is usually a good indicator of sample dilution and quality. The aim was to exclude outliers in both the female and male samples and to yield close to normal distribution in both (see **Figure 1** and **Suppl. Figure 1**).

The RNAseq TPM data, used in this study, are compositional data, referenced with respect to the whole transcript/library size. As such, they exist in simplex/Aitchison space[38] [39]. To allow derivation of quantitative relationships and obtain meaningful comparisons, correlations and distance measures, a transformation into Euclidian space was necessary and it was done by normalization using the geometric mean and log-ratio representation. Two more outliers were removed based on a preliminary PLS model, trained within the normal range of *GAPDH*, based on extreme outlier scores, **Suppl. Figure 2**. The remaining 242 samples were normalized by the geometric mean and log scaling (**Eqns. 1-2**) [38] [39], **Figure 1** and used in the further analyses.

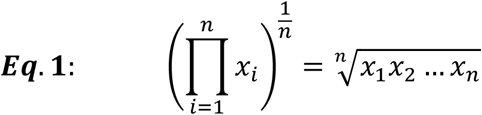

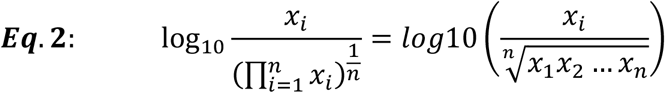

### Correlation analysis

The samples were analyzed in 3 blocks of relationships (HDACs, SIRTs, HATs to ion channels), (HDACs, SIRTs, HATs to TFs) and (TFs to ion channels), **Figure 1**. Pearson’s correlations **Eq. 3** (x, y representing the correlated variables, n – number of samples) were calculated for each one of these blocks and potential relationships visualized for the female and male samples.

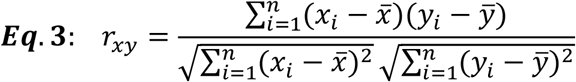

Clustering of the genes in **Figures 2, 3** was done using the clustergram function in the python library seaborn. The average and Euclidean were used for the method as distance metric. The algorithm used in the seaborn implementation is described in Bar-Joseph et al. [40]

**Figure 2:**
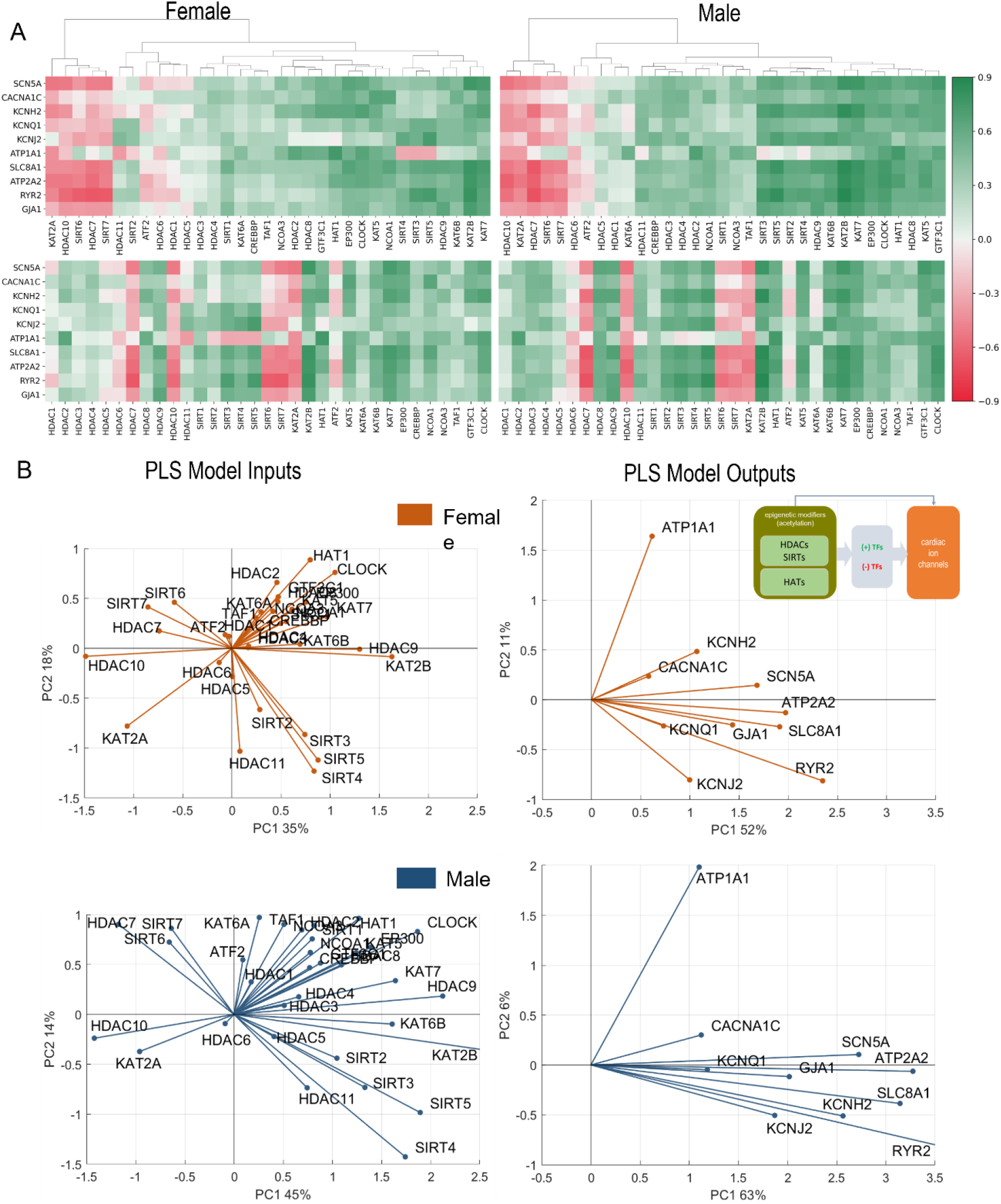
Analysis of links between histone acetylation modifiers (HDACs, SIRTs and HATs) and key cardiac ion channels based on transcription. A. Pearson’s correlation of ion channels with HDACs, SIRTs, and HATs for female (left) and male (right) LV heart samples. Positive/negative correlations are coded in green/red and shaded by their strength. The top correlation matrices are grouped using agglomerative clustering which generates the linkages. The unorganized correlation coefficients are on the bottom B. PLS regression models for female (orange) and male (blue) samples. Inset in B shows the models being investigated. The results are presented in biplots for the PLS model inputs (left) and model outputs (right). The biplots are projections of the model parameters onto the space of the first two latent variables of the constructed 4-latent-variable PLS models; shown are also the % variance explained for each latent variable. All biplots represent the average results from 1000 Monte Carlo PLS runs with random selection of training and testing samples (with the testing samples representing 10%). Proximity in angle between the lines for the model variables indicates co-regulation/similarity of action; the size of the lines signifies the importance of the variable in the shown projection. For example, *ATP2A2* is strongly influenced by the histone modifiers and is mainly represented in component 1 in the male and the female models (right panels). Positive co-regulation by *SIRT 3, 4* and *5* and negative co-regulation of *SIRT 6* and *7* on cardiac ion channels are strong in both the female and male models (left panels). See text for more interpretations.

**Figure 3:**
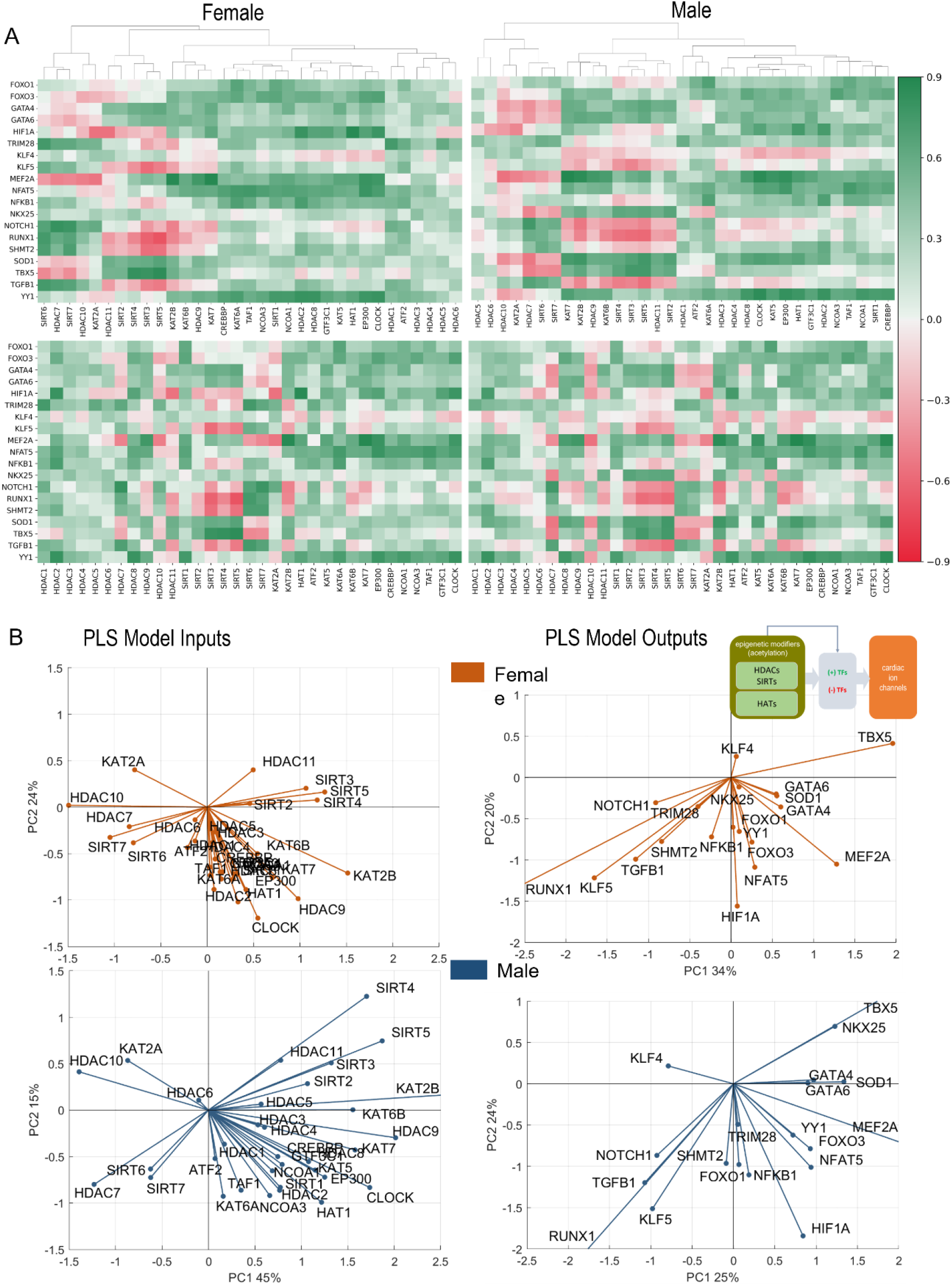
Analysis of links between histone acetylation modifiers (HDACs, SIRTs and HATs) and key cardiac TFs. A. Pearson’s correlation of TFs with HDACs, SIRTs, and HATs for female (top) and male (bottom) LV heart samples. Positive/negative correlations are coded in green/red and shaded by their strength. The linkages in the top clustering’s were created using agglomerative clustering. B. PLS regression models for female (orange) and male (blue) samples. Inset in B shows the models being investigated. Shown are biplots for the PLS models – constructed as described in Fig. 2.

### Partial Least Squares (PLS) analysis

Three types of PLS models were trained in MATLAB and Python on the female data and the male data separately. The only differences between the two platforms are the methods employed to execute PLS: MATLAB uses nonlinear iterative PLS (NIPALS) while the sklearn library uses the statistically-inspired modification of PLS (SIMPLS). Both methods produced similar results. When using PLS it is important to use the correct number of components. Once the model was trained, prediction was calculated using **Eq. 4**, where X and Y are the input and output variable vectors, B is the Beta-coefficient matrix calculated by PLS. Choosing the correct number of components can be found using cross-validation, percent variance explained, and prediction error sum of squares (PRESS). Leave-One-Out Cross Validation (LOOCV) was first used to find the number of components. Next PRESS was performed to confirm this finding. PRESS was calculated as per **Eq. 5**. The number of components settled on was 4, as it explained the most variance (over 70% for the various models) while keeping the model simple to prevent overfitting (see **Suppl. Table 1**).

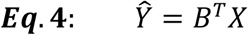

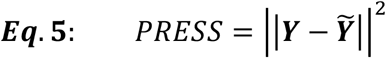

### Visualization by biplots

The results are presented in biplots for the PLS model inputs and model outputs. In all cases, the biplots represent the average results from 1000 Monte Carlo simulations (see below). The biplots are projections of the model parameters (PLS “loadings”) onto the space of the first two latent variables of the constructed 4-latent-variable PLS models. The biplots also indicate on the axes the % variance explained for each of the first two latent variables. For all models, the first two latent variables explained >50% of the variance. Proximity in angle between the lines for the model variables indicates co-regulation/similarity of action; the magnitude of the lines signifies the importance of the variable in the shown projection.

### Consideration of potential confounding factors and dimension reduction (UMAP)

Using phenotypic information about the GTEx samples, we considered several potential confounders to our female – male analysis, including age, BMI, quality index RIN, and total ischemic time, SMTSISCH. By feeding HDACs, SIRTs, HATs, TFs, Ion Channels into UMAP, a dimension-reduction visualization[41], the samples became embedded in a 2D space and were plotted and colored by the potential confounders. The only factor that appeared to systematically influence the clustering was the total ischemic time as shown in **Figure 5A**. Therefore, we performed a histogram matching for the female and male samples based on SMTSISCH, and built a “reduced” male model to compare to the female, as explained in the Results.

### Monte Carlo modeling and analysis of male and female differences

Monte Carlo Cross-Validation (MCCV) was used for all PLS models, typically 1000 runs were done with random assignment of training and testing samples (with 10% holdout for testing). This applies for the results shown in **Figures 2-4**. MCCV was also used to examine differences in the models trained on male and female samples, as shown in **Figure 5**. By randomly training n numbers of models a possible distribution of models was created. PLS was performed 1000 times on either the male or female samples. The male and female data was split into training and test sets with a random 10% holdout. After the split, PLS models were trained. The average B-matrix, scores, and loadings were calculated and plotted, and the data used for generation was stored for later use. Subsequently, two more tests were performed using the different inputs and outputs blocks. Each input and output block were analyzed to uncover features of regulation in finer detail.

**Figure 4:**
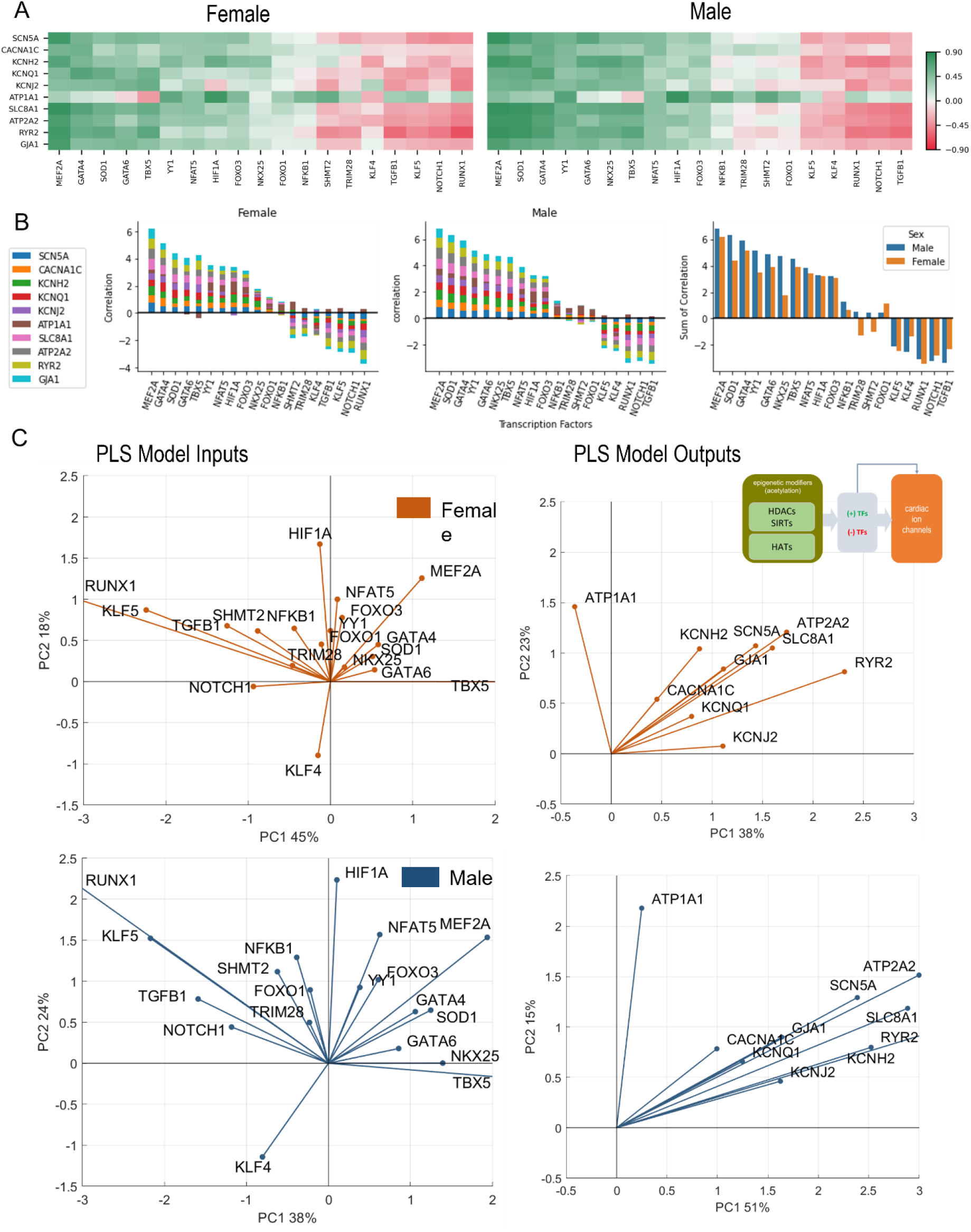
Analysis of links between cardiac TFs and key cardiac ion channels. A. Pearson’s correlation of key ion channels with cardiac TFs for female (left) and male (right) LV heart samples. The ranking of the TFs is done based on sum of correlations (from predominantly positive to predominantly negative) from left to right. These ranking yields slightly different order for female and male samples. B. Bar plots show the cumulative correlation of individual ion channels with each TF for female, male samples, and the final plot shows the overlaid impacts of TFs on ion channels for female and male samples to reveal any differences. C. PLS regression models trained with the TFs predicting ion channels for female (orange) and male (blue) samples. Inset in B shows the models being investigated. Shown are biplots for the PLS models – constructed as described in Fig. 2.

**Figure 5:**
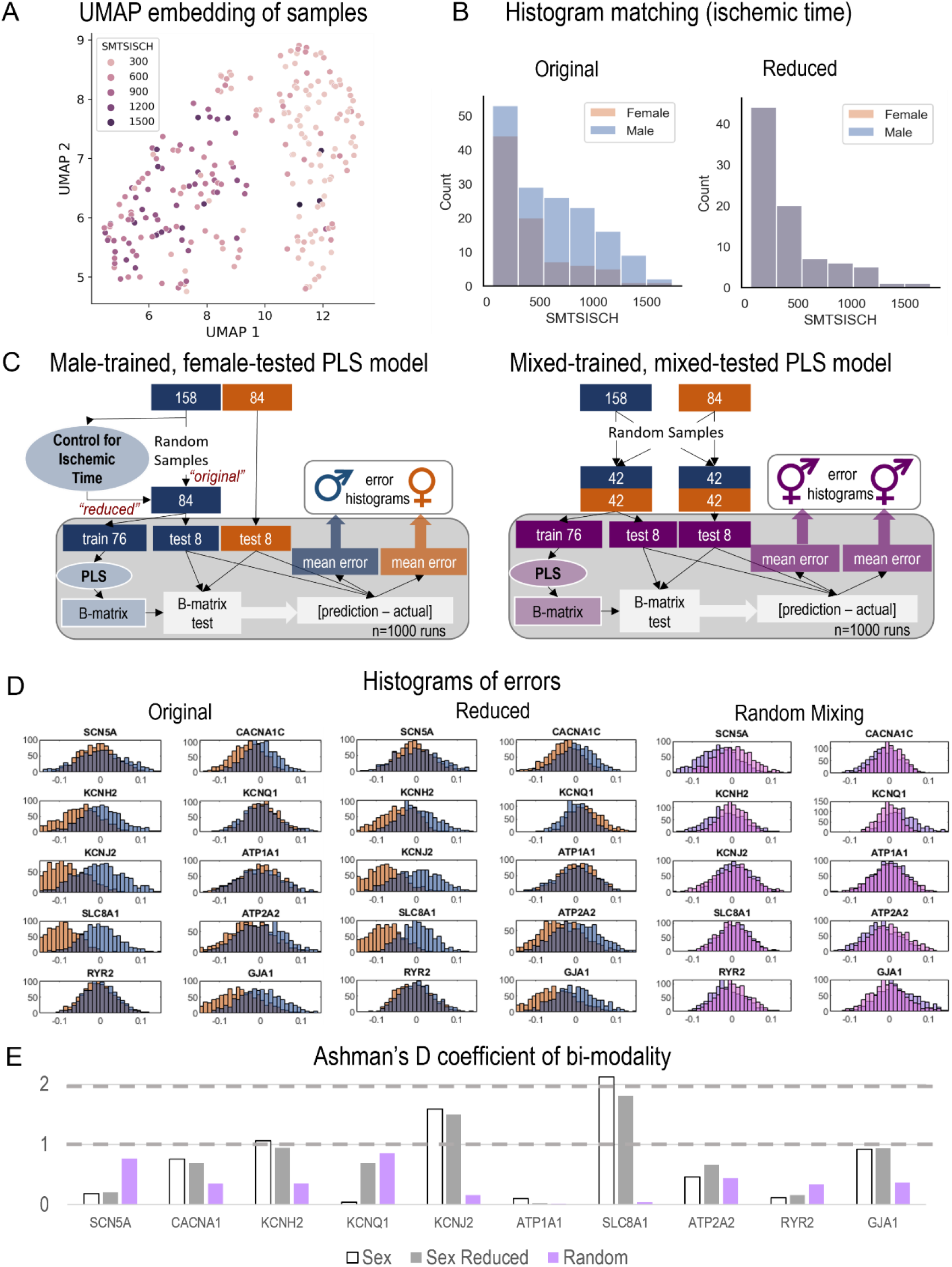
Monte Carlo (MC) simulations to examine the predictive power of male-trained PLS models when applied to female data. A. UMAP embedding (dimension reduction) was used to visualize the samples in 2D space, when considering all HDACs, SIRTs, HATs, TFs and Ion Channels. This representation showed strong role for the total ischemic time (SMTSISCH, colored) in the separation of samples. B. To eliminate the effects of SMTSISCH as a confounder in the sex-difference analysis, a histogram matching was performed - the histograms show the before and after reduction of the set of male samples to match the female samples based on total ischemic time. C. (Left) Details on the MC model runs and estimation of errors for male-trained, female-tested PLS models linking histone acetylation modifiers to cardiac ion channels in the LV. Two types of male models (matching the sample size of the female set, 84) were constructed – “original” with randomly selected 84 male samples and “reduced” (with histogram matching for SMTSISCH). (Right) Details on the construction of MC mixed models – trained on mixed data and tested on mixed female-male data. By mixing male and female, the model will have seen both samples and predict both well, assuming that the training samples represent the actual distribution of male and female samples. D. From the 1000 runs, the total errors were stored and displayed in histograms for the three types of models depicted in D, respectively. Note systematic shifts in errors in D from the zero-mean center. Left shift (to more negative error values) indicates underestimation of effects by the created PLS model for the respective variable. E. Ashman’s D coefficient (measure of bimodality, or how distinct the two populations of errors are) was calculated for each of the model results and displayed on the bar plot; higher Ashman’s D values indicate better separability.

Three tests to examine the differences between male and female samples were performed. Not all PLS model types were used to analyze the differences. The only relations examined were HDACS, SIRTs, HATs predicting ion channels. The first approach involved “male-trained, female tested models”. A model was created using only male samples for training (from the 158 male samples, 84 random samples were selected, of which 10% were used for holdout). With the 76 remaining male samples, a PLS model was trained. The beta matrix was obtained from this model, which is used to calculate the predictions. The errors were calculated as [*prediction – actual*] so that underpredictions would generate negative errors and over predictions - positive. Previous steps were repeated 1000 times with a new random hold out. For the next model a reduced data set was used using the same method. The dataset was reduced to control for the potential error introduced with ischemic time, as seen in **Figure 5A**. Once a bin width was chosen samples were removed randomly from the males within that bin until the number of samples in male and female matched. The third approach involved “mixed-trained, mixed-tested models”. After selecting 84 male samples, they were mixed with the female samples. This left two sets of 84 samples with equal numbers of male and female in each. This model was created to test if the relations are due to random sample bias. Both procedures are visualized in **Figure 5C**.

Beyond individual influence on ion channels, the potential effect on ion channel groupings were examined. These effects were calculated by taking the mean of the beta matrix coefficients for a desired ion channel group. The groupings analyzed were Depolarization (*SCN5A, CACNA1C, SLC8A1*), Repolarization (*KCNH2, KCNQ1, KCNJ2, ATP1A1*), Net (Depolarization – Repolarization), Resting membrane potential (*KCNJ2, ATP1A1*), Calcium Handling (*ATP2A2, RYR2, SLC8A1, CACNA1C*), and Coupling (*GJA1*). The coefficients used in these calculations were from two PLS models using all the samples (HDACs, SIRTs, HATs to Ion Channels and TFs to Ion Channels). From these Beta matrices the coefficients for a given group were averaged and then plotted using the seaborn library function barplot. The results of these calculations are seen in **Figure 6**.

**Figure 6:**
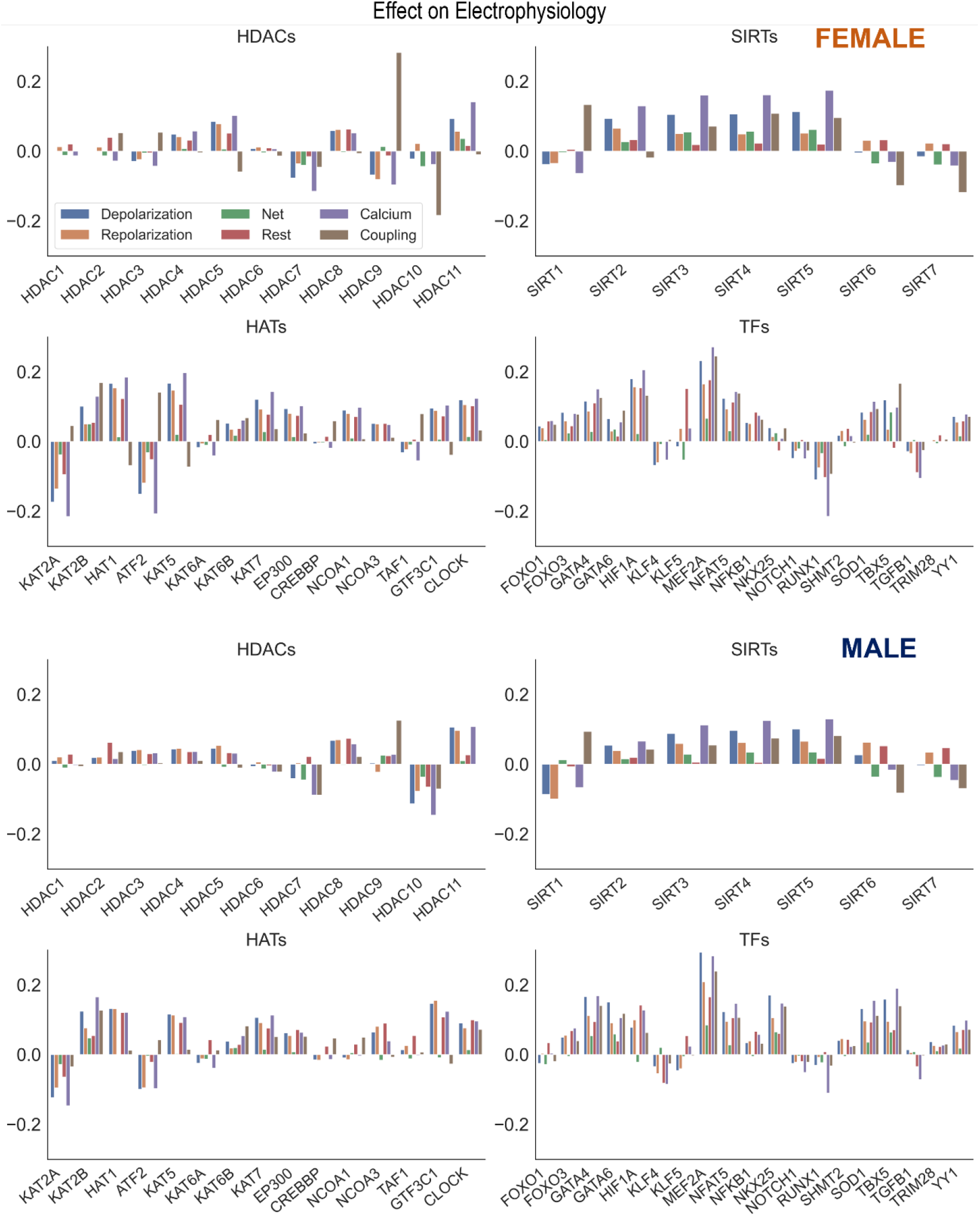
Control of key genes defining cardiac electrophysiology by HDACs, SIRTs, HATs, and TFs. Using the results from the PLS beta matrix, the effects of the HDACs, SIRTs, HATs, and TFs were summed to quantify the effect on the electrophysiology of cardiomyocytes. The 6 groupings examined were Depolarization (*SCN5A, CACNA1C, SLC8A1*), Repolarization (*KCNH2, KCNQ1, KCNJ2, ATP1A1*), Net (Depolarization – Repolarization), Resting (*KCNJ2, ATP1A1*), Calcium Handling (*ATP2A2, RYR2, SLC8A1, CACNA1C*), and Coupling (*GJA1*). This analysis was done for both female (top) and male (bottom) samples separately.

**Figure 7** was generated using the Sankey figure from the python library plotly. A new PLS model was trained in python using the 4 components. All samples were used to obtain the Beta Matrix used in the plot. Once the models were trained on male and female separately, the Beta Matrices were used for the coloring and generation of the plot. Since the scale of some coefficients in the Beta Matrix were large, it overshadowed the overall effect of most of the HDACs on TFs and TFs on Ion Channels. To mitigate this the MaxAbsScaler function from the library sklearn was used to scale the values of each gene between -1 and 1 without shifting the center, which preserves the sign of the effect. The weight and color of the lines were generated using the scaled data.

**Figure 7:**
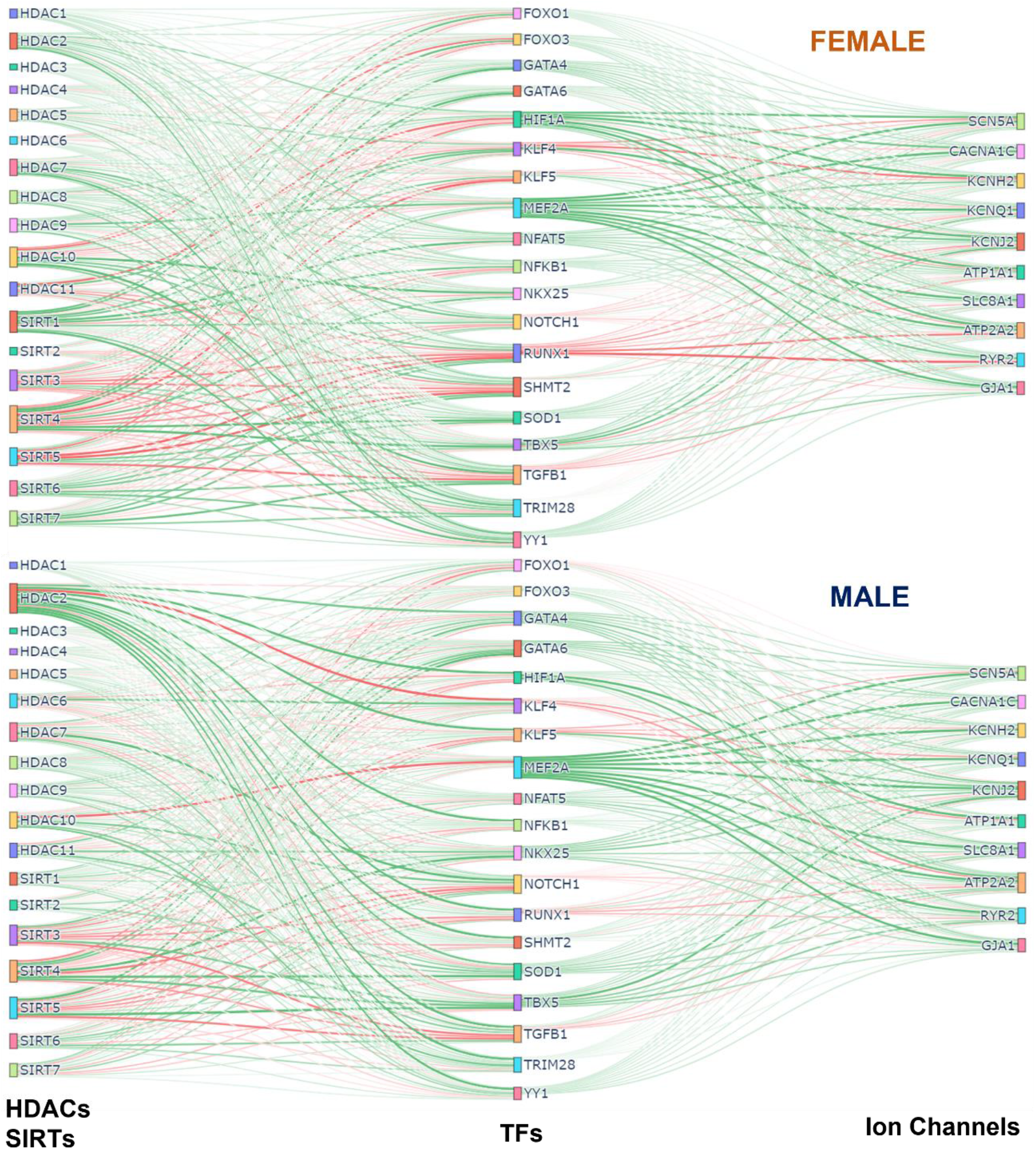
Integrated (Sankey) flow diagrams of how HDACs and SIRTs influence ion channels via cardiac TFs. PLS regression models (for female samples on top and male samples on bottom) were trained using HDACs and SIRTs to predict TFs and then using TFs to predict cardiac ion channels. The resulting B-matrix coefficients were utilized to create the plots. The width and color intensity of the lines correspond to the strength of the B-matrix coefficients, and the color distinguishes between positive (green) and negative (red) coefficients. Note that in some cases, double negative regulation (e.g. *SIRT3* -> *RUNX1* -> *RYR2*) results in positive input-output relationships.

### Statistical analysis

The Ashman’s D coefficient was used to examine if the two distributions of errors are part of the same distribution. For the two distributions to be separable the coefficient must be greater than 2. Ashman’s D coefficient is calculated using **Eq. 6**.[42]

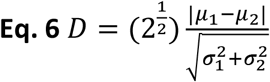

## Results

### Genes involved in the effects of histone acetylation modifiers on cardiac electrophysiology via key cardiac transcription factors

In this study various histone acetylation modifiers were considered: all HDACs, including class III, sirtuins, SIRTs; as well as key HATs, **Table 1**. From the 1500+ human transcription factors, previously a subset has been identified as “cardiac TFs”, of which we considered 19 that have been implicated in control of electrophysiological processes in the heart[31-33]. From the extended list of genes known to encode proteins relevant to cardiac electrophysiological processes in the LV, sometimes called the “rhythmonome”[14], we focused on the ten key genes controlling the depolarization – repolarization balance, the resting membrane potential, calcium handling and cell-cell coupling in the LV human heart, **Table 2**.

**Table 2.**
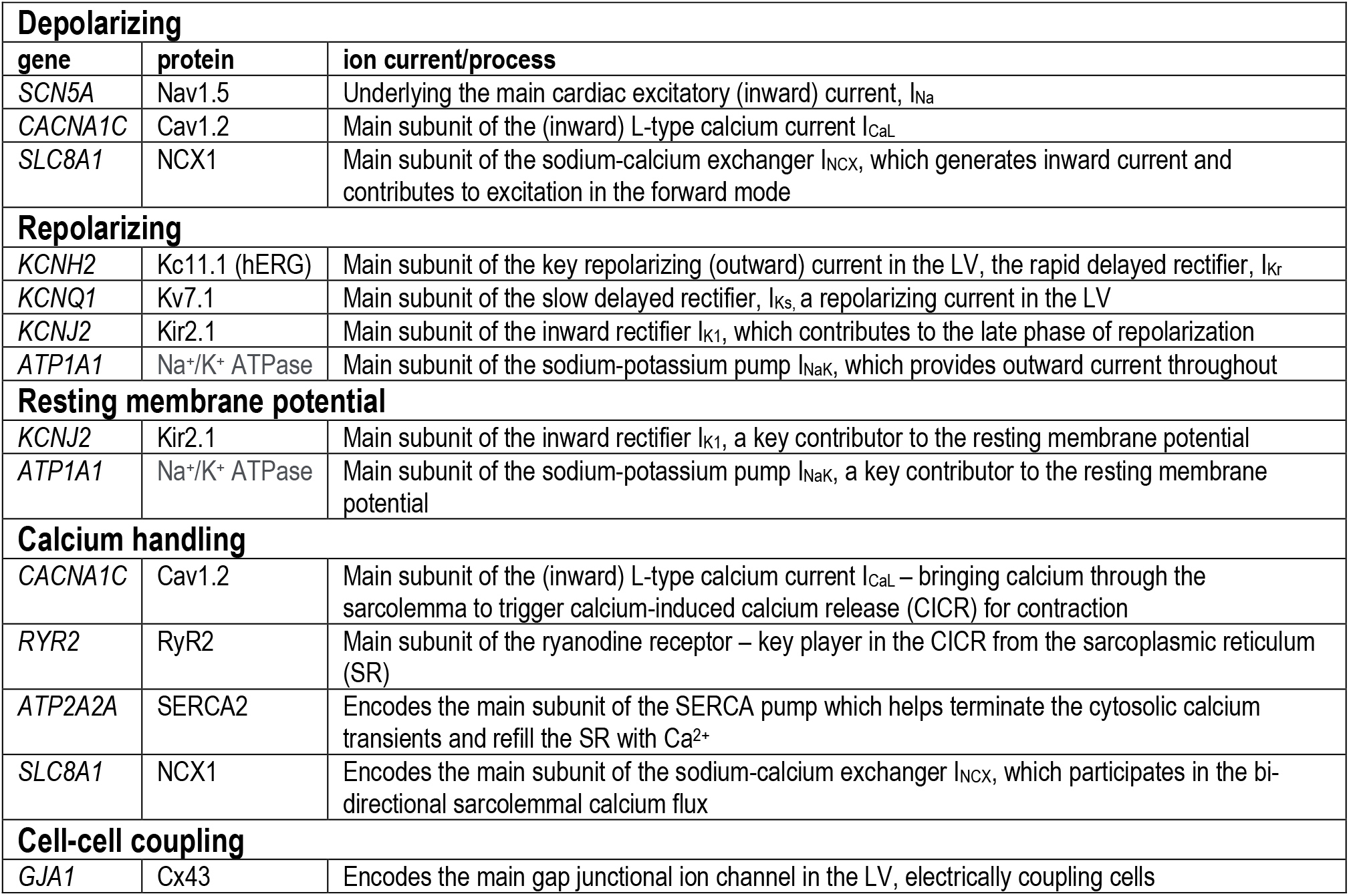
Genes, proteins and ion channels in the LV.

### Linking histone acetylation modifiers and key cardiac ion channels based on transcription

Hierarchical clustering of female and male correlation coefficients, **Figure 2A**, separates approximately 3 groups of negatively, weakly, and positively correlated epigenetic modifiers, i.e. HDACs and HATs, in relation to the select 10 cardiac ion channels, **Table2**. Many of these correlations are further corroborated by the results of PLS regression modeling on the transcriptomics data from female and male LV tissues, **Figure 2B**.

In female and male LVs, the strongest negative regulators of cardiac ion channels appear to be similar for the two sexes - *HDAC7, HDAC10, SIRT6, SIRT7* and *KAT2A*, based on correlations of gene expression. However, the strongest positive regulators are different in female and male LV. For male hearts, these positive regulators are mostly HATs (*GTF3C1, KAT5, HAT1, CLOCK* and *EP300*) plus *HDAC8*. In the female hearts, *SIRT3, SIRT4* and *SIRT5* along with *HDAC9* exhibit the strongest positive regulation, in addition to *KAT7* and *KAT2B*. The mitochondria-localized *SIRT3, SIRT4, SIRT5* act together, displaying very close (positive) relationships to the ion channel expression in both sexes. In contrast, the nucleus-localized *SIRT6* and *SIRT7* display strong negative correlations with cardiac ion channel expression in both sexes.

What is seen from the correlation analysis is that the key cardiac ion channels are largely co-regulated by these histone modifiers. An exception to this appears to be *ATP1A1*, which in many cases shows reverse correlation compared to the rest. PLS regression analysis results, presented as biplots in **Figure 2B**, provide a more granular view. For example, the co-regulation of ion channels by HDACs and HATs is more pronounced in the male LV compared to the female LV – tighter clustering and alignment seen in the PLS model output biplots. In both sexes, this key cardiac ion channel set is flanked by two genes regulating the resting membrane potential – *ATP1A1* and *KCNJ2*, with *ATP1A1* being orthogonally regulated compared to the rest. Such distinct pattern may have compensatory role – if the gene expression of all ion channels is upregulated or downregulated simultaneously, there may be undesired changes in the resting membrane potential that need to be countered. Genes involved in calcium handling (*RYR2, ATP2A2, SLC8A1*) are strongly regulated by histone acetylation modifiers in both sexes (see prominent representation lines in the output biplots). According to this PLS model, *SIRT2, SIRT3, SIRT4* and *SIRT5* act very closely together to exert positive regulation on the ion channels (exception being *ATP1A1*). Most of the HATs are co-aligned with *ATP1A1*, implying positive regulation of the Na/K pump to maintain the resting membrane potential. *KAT2A* is the exception in both sexes – it has distinct negative effect on all cardiac ion channels in both sexes along with *HDAC10* and to some degree *HDAC7, SIRT6* and *SIRT7*.

### Linking histone acetylation modifiers and key cardiac transcription factors

Most of the gene control by epigenetic modifiers is channeled through transcription factors that can act as positive or negative regulators. **Figure 3** shows our results from correlative analysis and clustering of such effects, **Figure 3A**, and from PLS regression modeling linking HDACs and HATs to cardiac TFs, **Figure 3B**. From the clustering analysis it is obvious that the effects on TFs are complex and the outcomes are not co-aligned like the ion channels. For some TFs, like *YY1*, most of the effects appear positive, especially for HAT effects on *YY1*. This means that histone acetylation by HATs likely yields increased expression of *YY1*. Similarly, for both sexes, the HATs result in positive regulation of one of the strongest cardiac TFs – *MEF2A*. Furthermore, *MEF2A* is negatively regulated by the same set of histone acetylation modifiers that appeared as the most prominent negative modulators of cardiac ion channel expression as per **Figure 2**: *HDAC7, HDAC10, SIRT6, SIRT7* and *KAT2A*. This reinforces the idea that the action of these epigenetic regulators on the ion channels is likely through potent TFs, as exemplified by *MEF2A*. Prior studies support some of these findings, e.g. *HDAC7* serving as a negative regulator of *MEF2A*[43],[31].

The PLS biplots reveal some differences between the female and male LVs. In the male LV, *NKX25* is co-aligned with *TBX5* and strongly influenced by the positive histone acetylation modifiers *SIRTs 2-5*. In contrast, the response of *NKX25* to histone acetylation modifiers in the female heart appears minimal. *TBX5* is strongly positively regulated by the mitochondrial SIRTs (*SIRT3-5*) and strongly negatively regulated by the nuclear SIRTs (*SIRT6* and *SIRT7*) in both sexes. Similar to *MEF2A*, likely *TBX5* plays an important role in mediating the effect of epigenetic modifiers on the ion channels. Other clustered response to histone acetylation is seen for *GATA4, GATA6* and *SOD1*, which show perfect co-regulation in the male LV along an axis similar but distinct from *MEF2A*. In the female LV, these additional TFs are closer to the responses for *MEF2A* yet smaller in magnitude. *GATA4, GATA6* and *SOD1* appear co-aligned with (likely positively regulated by) *KAT2B* and *KAT6B*. A distinct difference between the female and male model outputs is seen in *KAT2B* – this HAT is co-aligned (positive regulation) with *MEF2A* and *GATA4* in the female LV, while it is closer aligned with *GATA4* but not *MEF2A* in the male LV. Other positively regulated TFs, especially by the HATs are *HIF1A, NFAT5, FOXO1* and *FOXO3*. The responses of these four TFs are more closely aligned together in the female LV compared to the male. Many of the HATs, including *CLOCK, HAT1* and *EP300*, are clustered (act together) in their effects on the cardiac TFs and they are most closely aligned with *HIF1A, NFAT5, YY1, FOXO1* and *FOXO3*, implying positive regulation.

Distinct negative cluster of TFs is formed by *RUNX1, KLF5, TGFB1* and *NOTCH1*. For the female LV, these are joined by *TRIM28* and *SHMT2* and together they appear to present a counter-regulator for *TBX5*. Their expression may be positively regulated by *SIRT6, SIRT7* and *HDAC7* (negative regulators of ion channels as seen in **Figure 2**). In the male LV PLS model, *KLF4* appears as direct counter-regulator of *MEF2A*, presumably exerting negative effects on the ion channels. *HDAC10* and *KAT2A* seem to positively regulate *KLF4* in the male LV. *KLF4*’s role in the female model is less pronounced.

### Cardiac TFs and control of the gene expression of key cardiac ion channels

The set of 19 TFs was selected because of their known prominent role in regulation of genes underlying cardiac electrophysiology processes [31-33]. In **Figure 4**, we examine closely the role of these “cardiac TFs” on the ten ion channels of interest. As in **Figure 2** for links between the histone acetylation modifiers and the ion channels, here we also see pronounced co-regulation of ion channels by the TFs. This means that there are potent “positive TFs” that act to increase the expression of all or almost all key cardiac ion channels; there are also potent “negative TFs” that act to decrease the expression of all of almost all key cardiac ion channels. Among the positive TFs are *MEF2A, GATA4, GATA5, SOD1, TBX5, YY1*. Their extent of action varies in female and male LV. Perhaps the most striking sex difference is in the stronger role for *NKX25* on ion channel expression in the male LV compared to the female LV, **Figure 4B**-right panel. Additionally, some of the weaker TFs, like *TRIM28* and *SHMT2*, may show opposite effect on the regulation of the ion channels for the female and male LV. Based on the ranking of correlation in **Figure 4A-B** and based on the PLS biplots in **Figure 4C**, the ion channel co-regulation is also driven by a set of exclusively negative cardiac TFs. In this cluster are *RUNX1, NOTCH1, TGFB1, KLF5* and *KLF4* for both sexes, though their relative ranking is different for female and male LVs. Similar to the results in **Figure 2**, *ATP1A1* is orthogonally regulated (with respect to the rest of the cardiac ion channels) by many of the positive and negative TFs. This is clearly seen for *TBX5*, which exerts negative effect on *ATP1A1* yet acts as a strong positive regulator of the rest of the ion channels. Similarly, *RUNX1, TGFB1, NOTCH1* and *KLF5* exert positive effects on *ATP1A1* yet they act as strong negative regulators of the rest of the ion channels.

The PLS output biplots, **Figure 4C**, illustrate the co-regulation of cardiac ion channels by TFs. Again, similar to **Figure 2B**, this co-regulation is much stronger (tighter clustering) in the male LV. And the distinct orthogonal regulation of *ATP1A1* (compared to the rest of the ion channels) is seen in the female and male LV. *HIF1A* and *NFAT5* seem most aligned with *ATP1A1* (positive regulation) for both sexes. *TBX5, GATA6* (and *NKX25* for the male LV only) appear to be most aligned with *KCNJ2*. In both sexes, *MEF2A* plays a prominent positive role for the expression of cardiac ion channels, aligned with the cluster of ion channels. *KLF4* directly counters the role of *MEF2A*, especially in the male LV. The PLS input biplots also showcase the extent of clustering of the negative TFs, especially *RUNX1, KLF5, TGFB1, NOTCH1*.

### Can male-trained PLS models predict female relationships between histone acetylation modifiers and cardiac ion channels?

Using the genes of interest in this study, **Table 1**, we visualized all samples using a dimension-reduction approach, UMAP, **Figure 5A**. When color-coding various phenotypic variables associated with the samples, including age, BMI etc, we did not observe specific clustering, except for the UMAP plots when total ischemic time was displayed, **Figure 5A**. SMTSISCH is the total time for handling an LV tissue sample from the moment of opening the chest to the collection of tissue lysate for transcriptomics in GTEx. To eliminate ischemic time as a potential confounder in our analysis, we looked closely at the SMTSISCH histograms for female and male LV samples used. **Figure 5B**, left, shows that the larger data set for male LV did also include some longer ischemic times. We performed histogram-matching, by reducing the male dataset, eliminating samples so that the histograms of ischemic time for female and male samples became identical – see “reduced” in **Figure 5B**. The PLS biplots for the “original” male and female models along with the biplots for the new “reduced” male model are shown in **Suppl. Figures 3-5**.

We tested if purely male-trained PLS models of influences of HDACs and HATs on cardiac ion channels can accurately predict such influences for female LV samples. This was done using the full male set of 158 samples (**Figure 5C** – left, “original” model), the “reduced” male data set of 84 samples with histogram-matched ischemic time (**Figure 5C** – left, “reduced” model) and for a “mixed” model using equal number of female and male samples for training and for testing (**Figure 5C** – right). Monte Carlo simulations in all three cases yielded prediction errors for various outputs (ion channels) when the models were tested on male and female samples. Histograms of these errors are displayed in **Figure 5D** for the “original” and “reduced” male-trained models as well as for the “mixed” model. For good predictions, we expect the error histograms to be centered at zero and to be relatively tight. This is mostly the case for the mixed model, where the male and female error histograms overlap. The “original” and “reduced” male-trained models exhibit similar behavior for some of the ion channel genes, such as *SCN5A, KCNQ1, ATP1A1* and *RYR2*, implying that predicting the effects of HDACs and HATs on these genes may not be sex dependent. However, other results, especially for *KCNJ2, SLC8A1, KCNH2* and *GJA1* had error histograms for female predictions that were left shifted from zero, implying that the extent of regulation of these genes was under-estimated by the male-trained models. This is similar for the “original” and the “reduced” model (after ischemic time has been histogram-matched). Ashman’s D is a metric that can quantify the degree of bi-modality in a lumped distribution that may encompass two distinct populations. In **Figure 5E**, we computed Ashman’s D for the error histograms obtained in the three models. For the “mixed” model (training on equal mix of male and female samples), Ashman’s D coefficients remained low (<1) for all output variables. For the “original” male-trained model, Ashman’s D exceeded 1 for *KCNH2, KCNJ2* and *SLC8A1*. The “reduced” model showed similar trends to the “original” male-trained model, though slight decrease was seen in this metrics of bimodality. Ashman’s D was the highest for *SLC8A1*, implying that male-trained models may inaccurately predict the regulation of the Na/Ca2+ exchanger by HDACs and HATs in the female LV.

### Histone acetylation modifiers and net effects on cardiac electrophysiology

Our results showed strong co-regulation of the cardiac ion channels by HDACs, HATs (**Figure 2**) and by the cardiac TFs (**Figure 4**). We were interested to see the effects of these regulators on groupings of ion channels, as defined in **Table 2**: genes encoding for depolarizing, repolarizing ion channels, genes for ion channels and processes responsible for the maintenance of the resting membrane potential, for calcium handling or cell-cell coupling. The results of these effects, color-coded by groupings are displayed in **Figure 6** for the female (top) and the male (bottom) models. Of particular interest is if a certain perturbation may bias the net effect (positive for net depolarizing effect, negative – for net hyperpolarizing effect). As expected, most of the individual perturbations by HDACs, SIRTs, HATs and cardiac TFs leave minimal (close to zero) net effect. For the mitochondrial SIRTs and some of the positive TFs, there is a net depolarizing effect, based on the subset of ion channels considered here, possibly increasing excitability. Similarly, mitochondrial SIRTs and some of the HDACs (*HDAC 11*, and *HDAC5* for female only) plus some of the HATs (*KAT2B, HAT1, KAT5, KAT7, EP300, GTF3C1, CLOCK*) have prominent positive effect on genes related to calcium handling. Conversely, the strongest negative regulators of calcium handling appear to be *HDAC7, KAT2A, ATF2, SIRT1* (plus *HDAC9* for female LV and *HDAC10* for male LV). The impact of TFs on calcium handling genes directly follows from the results on “positive” and “negative” TFs as seen in the previous figures. The strongest positive regulators of calcium handling genes in the female LV are *MEF2A, HIF1A, GATA4, NFAT5* and *SOD1*, while in the male LV these are *MEF2A, TBX5, GATA4, NKX25* and *NFAT5*. The strongest negative regulators of calcium handling genes for both sexes appear to be *RUNX1, TGFB1* and *KLF4*.

### Sankey plots visualizing overall relationships between histone acetylation modifiers, transcription factors and cardiac ion channels

To visualize the overall effects of histone acetylation modifiers on cardiac ion channels via select transcription factors we used Sankey plots, based on the B-matrix coefficients from the PLS regression models for females (top) and males (bottom), **Figure 7**. The sign and the magnitude of the B-matrix coefficients determine the coloring and the thickness of the lines. Among the similarities between female and male, *MEF2A* appears to be the strongest positive TF of ion channels with a mix of positive and negative influences from HDACs and SIRTs. The strongest negative TF of ion channels, especially in the female LV, is *RUNX1*, which also gets mixed positive and negative influences from HDACs and SIRTs. For example, for both sexes, the mitochondrial NAD+ dependent sirtuins *SIRT3, SIRT4* and *SIRT5* downregulate *RUNX1*, thus exerting ultimately positive effect on the cardiac ion channels, while *HDAC7* and *HDAC10* positively influence *RUNX1*, thus having overall negative effect on ion channel expression. Other positive TFs of ion channels seen here are *GATA4, GATA6, SOD1, NFAT5, TBX5, YY1. HIF1A* is a significantly stronger positive regulator of ion channels in the female LV compared to the male, while *NKX25* is stronger positive TF of ion channels in the male hearts compared to female, in line with previous studies that have identified *NKX25* as a male-biased TF[8]. *HDAC2* also appears to play more prominent role in the male LV, regulating various cardiac TFs; *SIRT1* exerts stronger (positive) influences on cardiac TFs in the female LV. *SIRT3, 4* and *5* are prominent in both sexes, providing downregulation of the negative TFs and upregulation of the positive TFs, thus having an overall positive effect on cardiac ion channels.

## Discussion

In this study we considered the impact of epigenetic modifiers (HDACs and HATs) on the transcription of ion channels in the human heart based on the GTEx dataset. Histone acetylation status and the compactness of chromatin are known master regulators of gene expression. However, these epigenetic modifiers are much more studied and better understood for genes controlling growth or cell death[24, 44-46], often in the context of cancer. Their effect on bioelectricity-related genes is much less understood[31]. Cell excitation and calcium handling, as metabolically demanding processes, may be considered positively regulated during growth and hypertrophy, and downregulated during ischemia or other conditions when conservation of resources is needed. The prominent role played by sirtuins, especially the NAD+ dependent SIRT3-5 in regulating cardiac ion channels (**Figures 2-4** and **Figure 7**), reflects their role as linkers between metabolic state and post-translational modifications to control transcription according to available resources[47, 48].

It is not immediately clear if the classic views of HDAC activity leading to reduced transcription (of certain genes) due to compacted chromatin, and HATs activity leading to enhanced transcription due to more accessible chromatin are applicable in this case. To explore these regulations, we developed models linking the gene expression of histone acetylation modifiers to the expression of cardiac ion channels via select cardiac transcription factors. As the GTEx dataset is still with a limited number of samples (a total of 242 analyzed here), the applicability of some machine learning methods that thrive on large datasets is limited. We chose to use partial least-squares regression (PLS-R) in this study to quantify these relationships and to explore potential sex differences in them. PLS-R was developed by Svante Wold[34-36] to deal with noisy and collinear variables. It is most closely related to Principal Component Regression (PCR). Both methods remove the multicollinearity between samples by projecting them into a latent space. As PCR maximizes the variance explained for the new latent variable, it may overlook the possibility that the latent variable may not explain the predicted variable. This leads to using many more components then needed to predict the outputs. PLS-R, used here, on the other hand maximizes the covariance between the input and output variables, and therefore maximizes the predictability of the inputs-to-outputs links. This connection between the inputs and outputs makes PLS-R a desirable method for examining RNA-seq data for specific relationships. This method is interpretable, and it holds causal power (beyond simple correlation) if large portion of the variance in the experimental data can be explained by the model[34, 36, 49].

In **Figures 2-4** we show how the histone acetylation modifiers may link to cardiac ion channels via cardiac TFs. It is important that some of the key results could be corroborated by prior research, mostly done in rodents, where transgenic models can be used to delete genes of interest and dissect functional impact. One of the main results in this study is the tight co-regulation of genes encoding cardiac ion channels by the histone acetylation modifiers and by the cardiac TFs. This is seen as clustering and co-alignment of the ion channel genes in the PLS output biplots in **Figure 2** and **Figure 4**, with the male hearts exhibiting tighter clusters compared to the female. These results are in agreement with a recent report[50], showing highly coordinated transcriptional control of cardiac ion channels in the ventricles, but not in the atria. This may be due, at least in part, to the reported synergistic action of positive TFs in the heart[32, 51] – collaboration between *MEF2A, GATA4, TBX5, NKX25* and the HAT *EP300*, identified based on chromatin co-occupancy through Chip-seq analysis of mouse heart tissue. Furthermore, *GATA4* has been linked to chromatin acetylation state (H3K27ac) in facilitating gene expression in the (mouse) heart. In the current report, using human RNAseq data, we see similar synergism between these strong positive TFs, including *GATA4*, in the regulation of cardiac ion channels. Furthermore, strong negative TFs mediating the effect of HDACs/SIRTs/HATs on cardiac ion channels, such as identified here *RUNX1*, have been reported in recent studies as well[52, 53]. Specifically, these reports showed adverse cardiac effects, including suppressed calcium handling by *RUNX1* after myocardial infarction[52, 54], in line with our results. Another strong negative TF shown here is *KLF5*, closely co-aligned with *RUNX1. KLF5* has recently been identified as a key negative regulator in ischemic cardiomyopathy in mouse and human hearts[55, 56] and has been suggested as a potential therapeutic target for drug development.

A recent study[14] combined meta analysis of human RNAseq datasets, computational modeling of cardiac electrophysiology and *in vitro* experiments with human induced pluripotent stem cell derived cardiomyocytes (iPSC-CMs) to show a strong correlation between *CACNA1C* and *KCNH2* and better stability of cardiac electrical activity (robustness and protection against arrhythmias) for high co-expression/co-regulation of these two genes. In our results, *CACNA1* and *KCNH2* are perfectly co-aligned/co-regulated by HDACs/SIRTs (**Figure 2B**) and by cardiac TFs (**Figure 4C**) in the female LV, and less so in the male LV. If these sex differences are confirmed in further experiments in human cardiac cells and tissue, they could partially contribute to higher resistance to arrhythmias in female hearts.

The co-regulation of ion channels controlling depolarization and repolarization implies that the HDACs/SIRTs/HATs and cardiac TFs are likely to leave mostly net zero effect in terms of excess depolarization or excess hyperpolarization in the heart, as seen in **Figure 6**. This is important for overall electrical stability. It may also explain the relatively low cardiotoxicity of HDAC inhibitors [27]. Even if the effects of HDACs/SIRTs/HATs and cardiac TFs do not create net depolarization/hyperpolarization, a concerted significant upregulation of ion channel genes can lower cell/tissue impedance; conversely, a concerted downregulation of ion channels will increase tissue impedance. The former will make it harder to excite or for an extra beat to propagate; while the latter may increase the risk for an aberrant beat to propagate. These more subtle effects of HDACs/SIRTs/HATs modulating cardiac ion channels, in tandem, may impact how epigenetic modulation alters arrhythmia propensity[28-31].

A distinct orthogonal regulation of *ATP1A1* (with respect to the rest of the cardiac ion channels) was seen by the HDACs/SIRTs/HATs and by the cardiac TFs, **Figure 2** and **Figure 4**. This may be a way to compensate for altered impedance (see above), as *ATP1A1* encodes for the Na/K pump, a major determinant of the resting membrane potential. It has been shown previously that higher level of excitatory activity and higher calcium release in neurons regulate *ATP1A1* transcription towards such compensatory effect[57]. *HIF1A* has been identified as a TF closely linked to *ATP1A1* regulation[57], and our data corroborate that, as seen in **Figure 4C** for both female and male.

Sex differences in transcriptional processes related to membrane ion transport, excitability and interaction with drugs[7], as well as in chromatin regulation and epigenetic influences[21],[58] have been previously reported for various organs. In the heart, the consequences of such sex differences may be of prime importance when developing and testing new drug compounds, including HDAC inhibitors. Recently, cardiac simulations showed that models integrating male-based data underestimated the risk for lethal arrhythmias, such as Torsade de Pointes, in females[18]. To aid the comprehensive preclinical analysis of drug action, transcriptional and other relevant data need to be viewed in a stratified manner, with considerations of the target demographics, including sex[6]. In the current study, we show several distinct sex-dependent patterns of regulation of histone modifiers and their action on cardiac ion channels. Specifically, we showed that male-trained PLS models may underestimate the effects of HDACs/SIRTs/HATs on certain ion channel related genes, including *SLC8A1, KCNJ2* and *KCNH2*, **Figure 5**. Other insights from the PLS analysis, including sex differences in the tightness of cardiac ion channel co-regulation, the more prominent role of *NKX25* the male LV, and stronger effects of *RUNX1* in the female LV, need to be confirmed as larger human datasets become available, as well as tested explicitly experimentally.

The analysis provided here has certain limitations – we worked only with gene expression data, although understanding of the complex action of epigenetic modulators can be enhanced through additional chromatin accessibility assays. While PLS-R may suggest input-output causality, ultimately such relationships need to be tested experimentally. The data provided here corroborates some previous relationships, but also shows new interactions that need to be tested experimentally. Larger datasets are likely to help better model development and to increase the confidence in model predictions. Our current analysis was done on bulk tissue RNAseq data and cannot differentiate if sex differences may be due to differences in cell type composition. Recent studies have suggested sex-distinct cell type composition, including a larger fraction of myocytes in the female LV compared to the male LV[59], which may explain certain sex differences in transcriptional regulation. Ideally, single-cell or spatial transcriptomics data[60], combined with the framework developed here, can improve the mechanistic understanding of the action of histone acetylation modifiers in the heart. Moreover, a small selection of genes was analyzed (number of genes smaller than the number of samples) in this study, however, PLS can further be utilized with the full transcriptome. For cases where much longer list of genes is considered – longer than the number of samples, a “sparse PLS” can be used as a dimension-reduction and a parameter selection tool[61].

In conclusion, we present a framework that aims to help the better understanding of how epigenetic modifiers may alter cardiac electrophysiology based on RNAseq data from human hearts. With the rapid growth and availability of such data it is possible to refine computational models to reliably predict responses to various histone modifiers in a stratified, personalized manner.

## Acknowledgments

This work was supported in part by grants from the National Science Foundation, EFRI 1830941, and the National Institutes of Health, R01HL144157, to EE.

## Author Contributions

MP, AH and EE designed the study. AH accessed the GTEx database, extracted the data, performed initial filtering and checks. MP and EE constructed the models and interpreted the results. MP wrote all codes related to the project, performed the analysis, and created the figures. EE supervised the project. MP and EE wrote the paper with input from AH. All authors reviewed and revised the manuscript.

## Code availability

The Python and Matlab scripts used to generate the results in this study are made available through github: https://github.com/pressm/PLSGTex.

