## Supplement for "Sex-dependent transcriptional control of cardiac electrophysiology by histone acetylation modifiers based on the GTEx database"

for

**Supplemental Table 1**. Percent variance explained in the PLS models. Three models were trained on male and female data. The table contains the average percent variance, explained from 1000 runs of the models (using Monte Carlo) as the PLS components increase from 1 to 4.

| | | Male | | | | Female | | | |
|---|---|---|---|---|---|---|---|---|---|
| HDACs, SIRTs, HATs --> Ion Channels | n components | 1 | 2 | 3 | 4 | 1 | 2 | 3 | 4 |
| | X | 45.49% | 13.76% | 12.78% | 4.11% | 34.99% | 17.91% | 21.27% | 3.36% |
| | Y | 63.12% | 5.75% | 2.92% | 3.32% | 52.38% | 11.13% | 1.61% | 6.27% |
| HDACs, SIRTs, HATs --> TFs | X | 44.77% | 14.82% | 12.94% | 3.10% | 29.28% | 24.09% | 21.10% | 3.18% |
| | Y | 24.7% | 24.48% | 3.60% | 6.27% | 33.99% | 20.34% | 7.35% | 4.60% |
| TFs --> Ion Channels | X | 38.1% | 24.01% | 7.26% | 6.10% | 44.81% | 18.42% | 11.37% | 6.11% |
| | Y | 51.37% | 15.07% | 2.06% | 1.75% | 38.49% | 22.71% | 3.45% | 2.46% |

**Supplemental Figure 1. Pre-filtering of the data based on GAPDH TPMs.** Any samples that were too high or low in GAPDH (outside of 500 to 2500 TPMs) were removed as potential outliers.

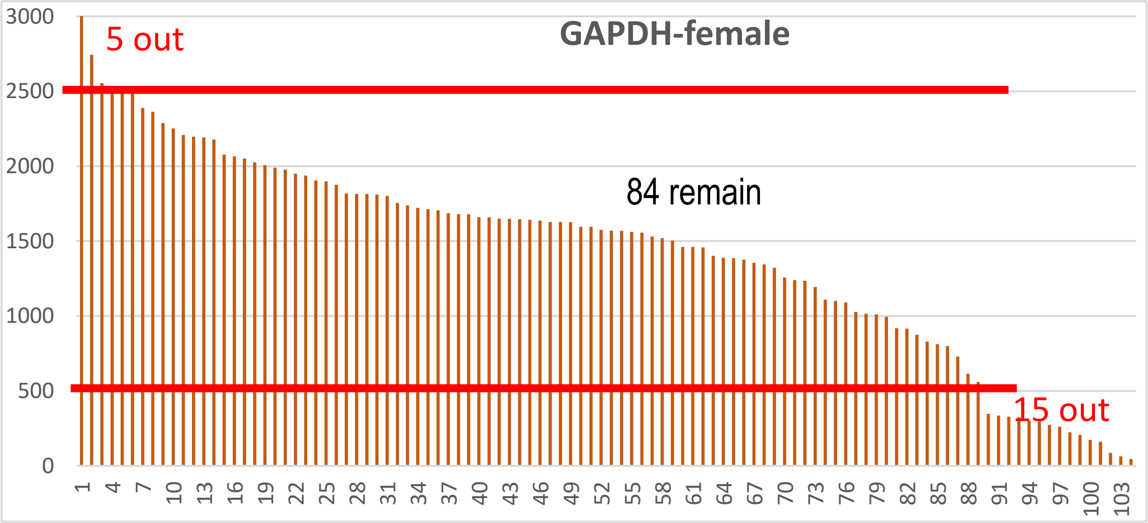


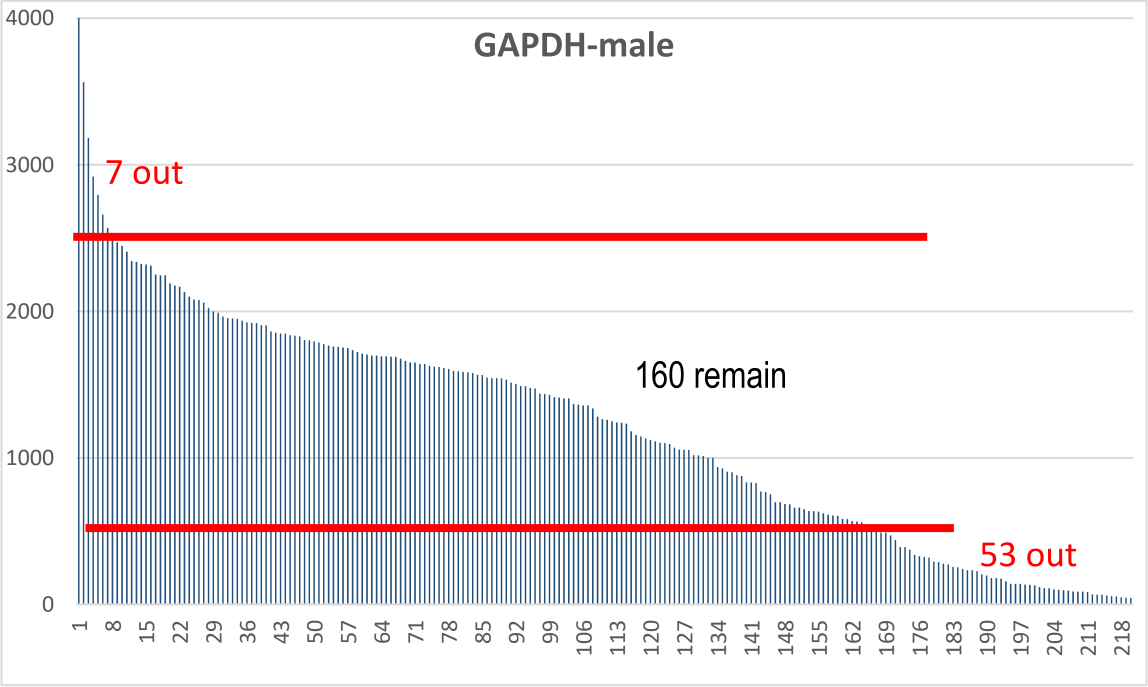

**Supplemental Figure 2. Outlier sample detection using PLS score plots.** After training the PLS models, the score plots can be used to find outliers. Outliers may cause the model to skew in a way that does not represent the general population. By plotting the X and Y scores of each of the 4 model components, a close to linear relationship is expected. Samples 113 and 131 (GTEX-1JJE9, GTEX-1PPH7) were removed from further analysis as outliers (see panel A).

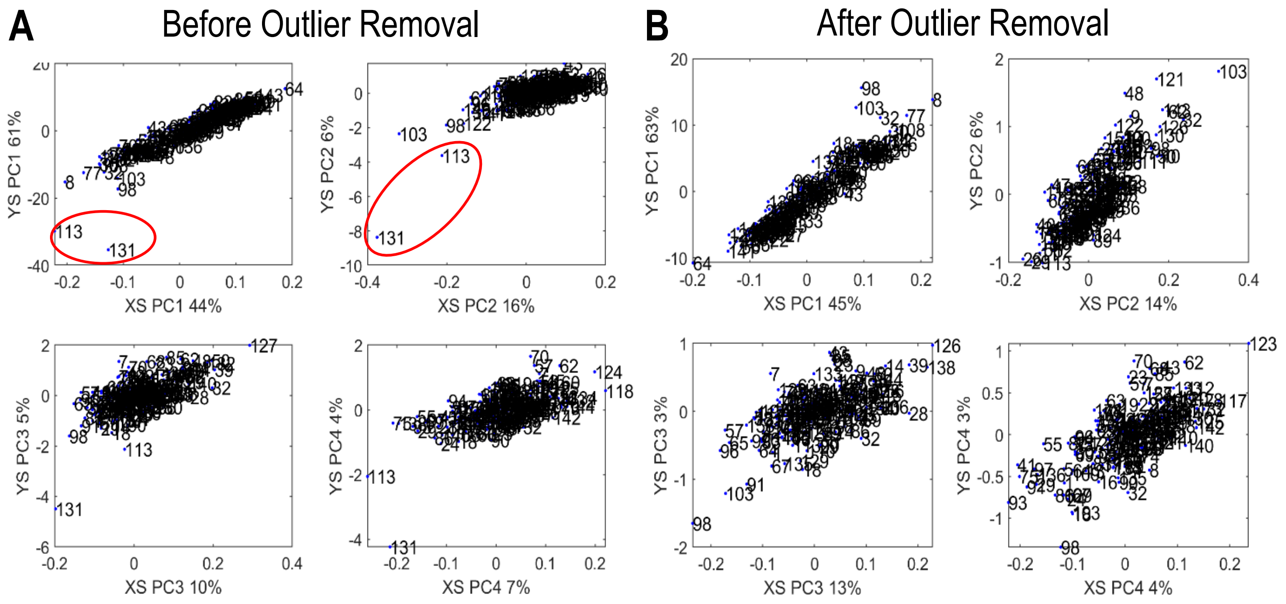

**Supplemental Figure 3. Biplots for the "original" PLS1 female and male models (as in Figure 2) along with the "reduced" male model obtained after histogram-matching of ischemic time (see Figure 5B).**

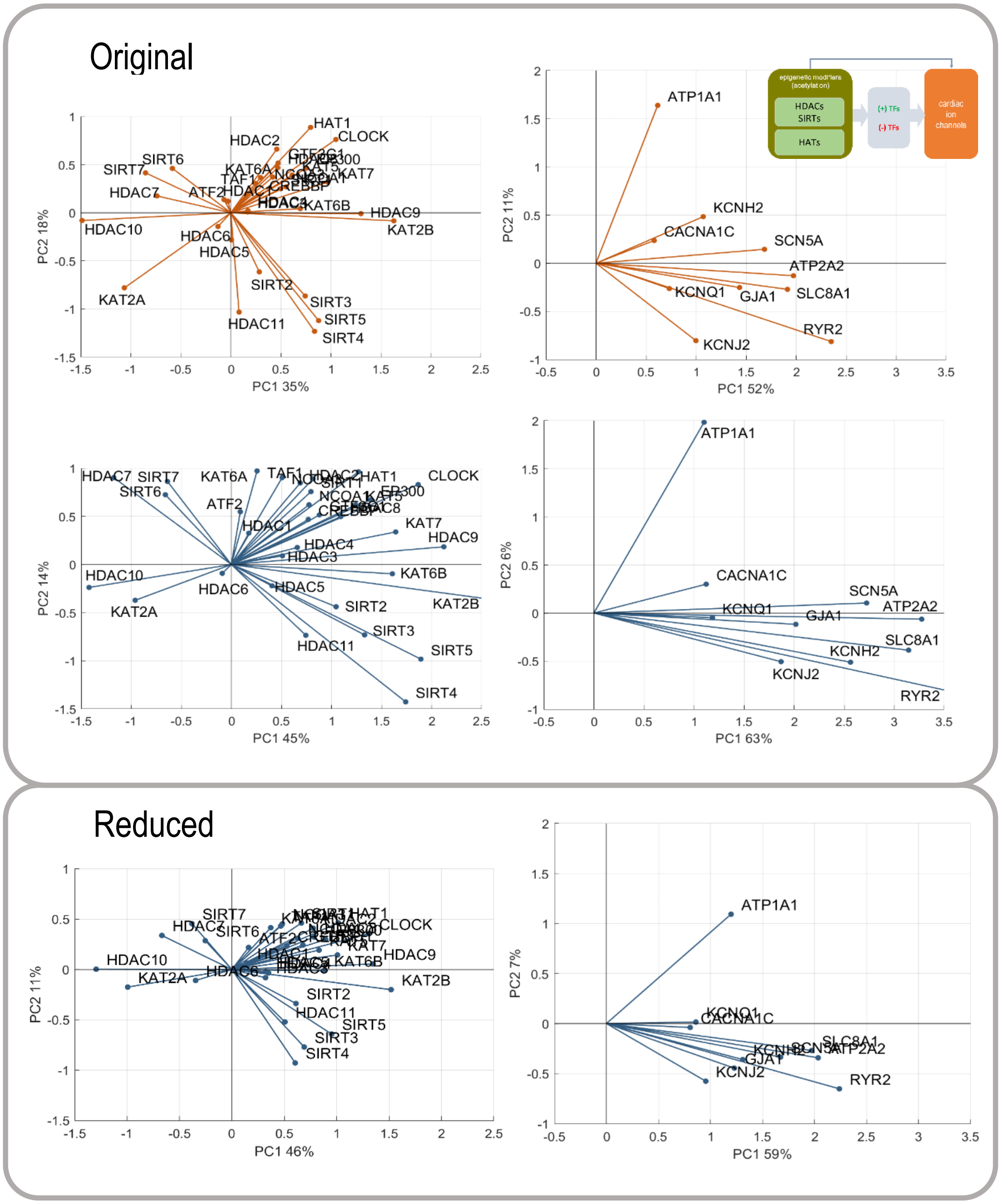

**Supplemental Figure 4. Biplots for the “original” PLS2 female and male models (as in Figure 3) along with the “reduced” male model obtained after histogram-matching of ischemic time (see Figure 5B).**

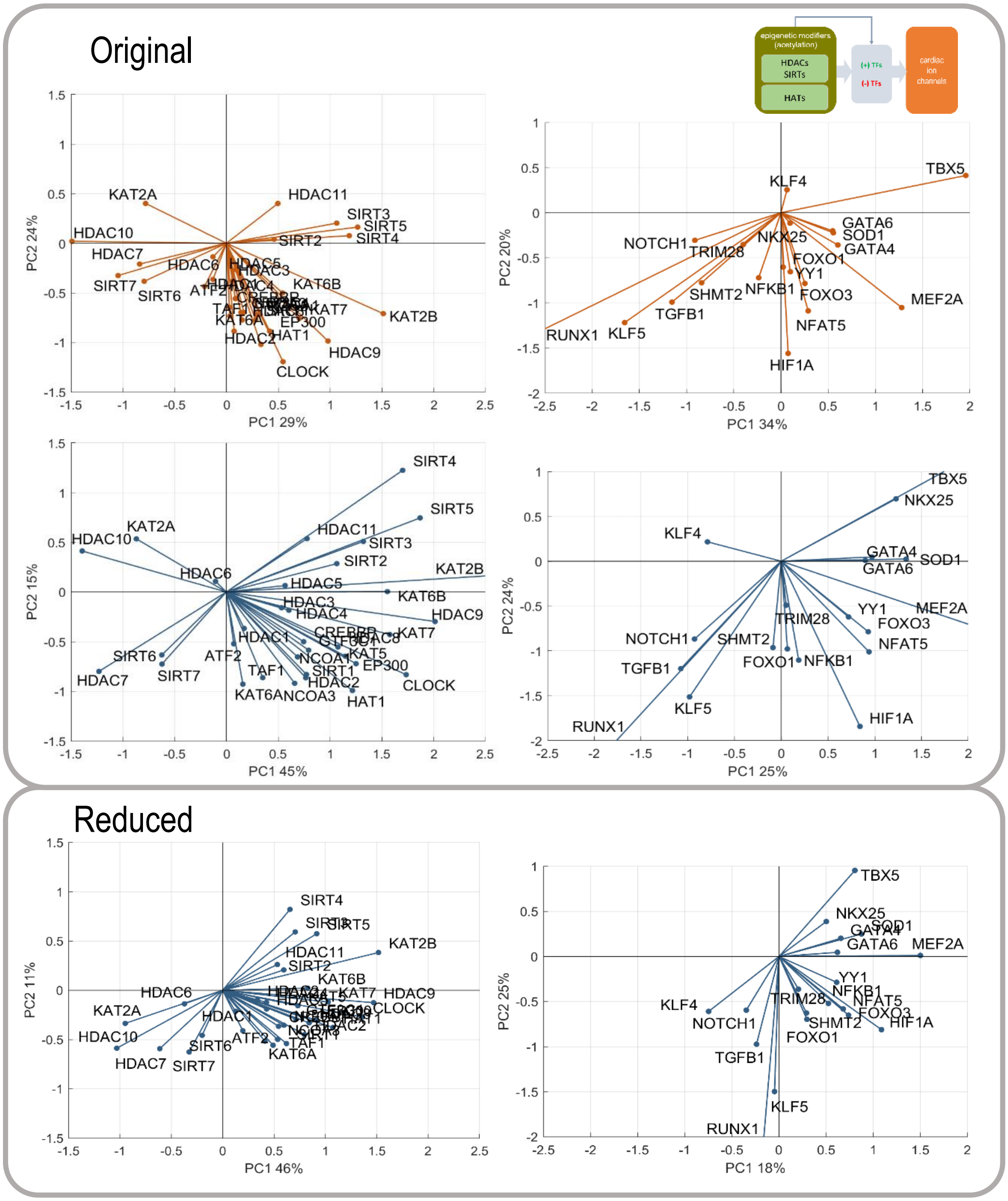

**Supplemental Figure 5. Biplots for the "original" PLS3 female and male models (as in Figure 4) along with the "reduced" male model obtained after histogram-matching of ischemic time (see Figure 5B).**

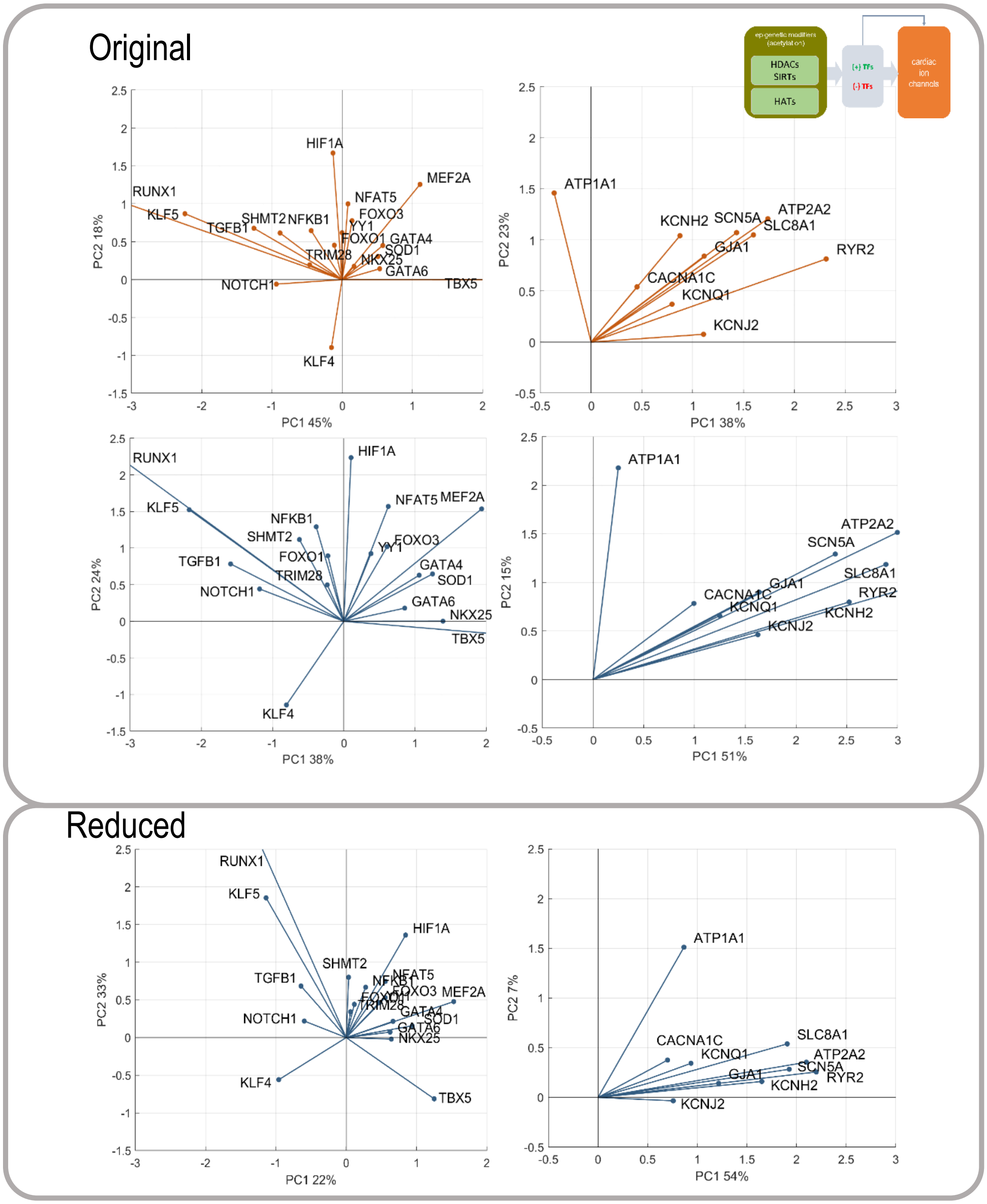
